# Does being a grateful person increase one’s level of well-being? The mediating role of indebtedness

**DOI:** 10.1101/344440

**Authors:** G Bernabé-Valero, C Moret-Tatay, T Navarro-Sancho

## Abstract

In this work, we define gratitude, paying attention to interpersonal gratitude and its relationship with dispositional debt. We examined the disposition to feel indebted through analysis of convergence and divergence, exploratory and confirmatory analysis of the most used measurement instrument. The Revised Indebtedness Scale depicted a four factor solution interrelated with a high consistency of content, which allows their labeling and describing. To do this, two samples of university students were selected; one of the sample sizes had 229 Spanish participants and the other 200 participants. Subsequently, a mediation model was tested in which the “Self-sufficiency and discomfort in receiving help” factor mediates the relationship between interpersonal gratitude and the “Positive relations with others” dimension of the Wellbeing scale. The results are discussed in relation to the need for conceptual definition of the constructs in Positive Psychology.

## Introduction

The importance of gratitude in the current field of psychology is indisputable, since gratitude has been identified as one of the most relevant indicators of a good quality of life (Solom, Watkins, McCurrach & Scheibe, 2016). It has been found that the gratitude trait is correlated with a large number of well-being variables (for example, Emmons & Misrha, 2011; García-Méndez, Serra-Desfilis, Márquez-Barradas, & Bernabé-Valero, 2014; McCullough, Emmons & Tsang, 2002; Watkins, Woodward, Stone & Kolts, 2003; Wood, Joseph & Maltby, 2008). On the other hand, several experimental studies have consistently found that fostering gratitude improves subjective well-being (for example. Emmons & McCullough, 2003; Lyubomirsky, Dickerhoof, Boehm, & Sheldon, 2011; Seligman, Steen, Park, & Peterson, 2005; Watkins, Uhder & Pichinevskiy, 2015; Watkins, 2014; Wood, Froh & Geraghty, 2010).

A large body of research illustrates and sheds light on this relationship between gratitude and well-being. For example, Emmons & Stern (2013) affirm that working on and promoting gratitude can be beneficial for our mental health, as it provides us with positive experiences such as well-being, happiness, positive affect and prosocial behaviors. It also acts as a protective factor against negative emotions. The authors, Ramírez, Ortega, Chamorro & Colmenero (2014) applied a positive psychology intervention, based on autobiographical memory, forgiveness and gratitude, with the aim of improving the quality of life and subjective well-being in older adults. The results of this study showed that gratitude intervention had a positive effect on the general well-being of these people. Killen & Macaskill (2014) used a gratitude diary in which participants had to write down three good things for which they felt grateful. They demonstrated that this exercise significantly increased psychological well-being; specifically, it improved the hedonic and eudaimonic wellbeing while reducing the levels of stress and also these results were maintained in 30-day follow-ups. In a study carried out by Watkins et al. (2015), it was concluded that there is a causal link between gratitude and well-being in the context of positive gratitude interventions. In this study, participants who wrote down three things for which they felt grateful each day over the course of a week, compared to those who wrote down things that increased their pride or those in a memory placebo condition, experienced greater subjective well-being until five weeks after the intervention. More recently, Krejtz, Nezlek, Michnicka, Holas & Rusanowska (2016), carried out a study where they examined the existing relationship between gratitude and well-being, and they discovered that gratitude and well-being were positively related.

In sum, these three variables are more related to interpersonal gratitude. Cynicism could have a detrimental effect on gratitude because if one believes that the benefactor provided a benefit for hidden or ambiguous reasons, the beneficiary could interpret that there is an expectation of future favors, so that gratitude would be less likely to occur. (Watkins, Scheer, Ovnicek & Kolts, 2006). Regarding narcissism, it has been found that it could inhibit gratitude and, conversely, humility should encourage gratitude (Watkins, 2014; Emmons & Crumpler, 2000; McCullough, Kilpatrick, Emmons & Larson, 2001; Roberts, 2004), since a person who believes he deserves a particular gift finds it more difficult to recognize the benevolence of the donor (McCullough et al., 2001). A key aspect for gratitude according to Seligman (2011) is that the person may feel unworthy of the gifts received, which corresponds to an attitude of humility. One of the variables that has been most related to the inhibition of the emergence of interpersonal gratitude is indebtedness (Watkins, 2014), a main aim of the present work.

We also consider that the willingness to feel indebted to the person who provides a gift would be one of the key points to inhibit gratitude and therefore, that this could have less effects on welfare. Several studies have found that the gratitude and indebtedness traits are negatively correlated (Elster et al., 2005; Van Gelder et al., 2007) in such a way that those people who showed a greater predisposition to feel indebted when receiving a gift, in turn showed a lower predisposition to feel gratitude. Although some theorizations have attempted to equate gratitude with indebtedness, arguing that returning a benefactor a favor could occur due to both indebtedness and gratitude, the researchers have attempted to define the differences between both concepts. In this way, it has been argued that indebtedness is accompanied by negative emotions, linked to discomfort (Greenberg, 1980), whereas gratitude is an emotion with positive valence. (Lazarus & Lazarus, 1994; Mayer, Salovey, Gomberg-Kaufman & Blainey, 1991). Based on his theory of expansion and construction of positive emotions, Fredrickson (2004) proposes that when experiencing gratitude as a pleasant emotion and debt as aversive, only gratitude could lead to a comprehensive and creative thought on how to return a favor. Secondly, indebtedness is associated with motives of avoidance, while gratitude is associated with rapprochement and prosocial motivations (Gray, Emmons & Morrison, 2001). Therefore, when it is understood that the intentions of the benefactor are benevolent, a greater response of gratitude occurs (Tsang, 2006), whereas the reasons or expectations of reciprocity of the benefactor would lead to an indebtedness response (Watkins et al., 2006). Thirdly, indebtedness arises from the reciprocity rule, while gratitude can go above and beyond the “eye for an eye” mentality (Fisher, 1983, Greenberg, 1980). Hence, gratitude might imply the perception and recognition of a gift, which is conceived as valuable, for which a positive emotion of connection with the donor is experienced, whilst indebtedness would imply a state of obligation to return a benefit to another, with an emotional experience of arousal and discomfort (Greenberg, 1980).

In previous studies, there have been few works carried out that measure the extent to which indebtedness diminishes the experience of gratitude. However, recent research carried out by Solom et al. (2016) aimed to study the “thieves of thankfulness”, the four supposed inhibitors of gratitude: - narcissism, cynicism, materialism / envy and indebtedness. The study concluded that narcissism, cynicism and materialism inhibit gratitude, with narcissism and cynicism being the main “thieves of gratitude”. With regards to indebtedness, it was predicted that the gratitude trait decreases with time (measured by the GRAT-S), but due to the inconsistency of the relationships between indebtedness and gratitude, it is difficult to draw clear conclusions (Solom et al., 2016). On the other hand, Mathews and Green, (2010), investigated the role played by self-focused attention in the development of the indebtedness feeling and in the decrease of the gratitude response. They found that when receiving a gift, people with a high level of self-focus would be more likely to experience indebtedness and, therefore, diminished gratitude.

Thus, it seems clear that upon receiving a gift, either a feeling of gratitude or a feeling of indebtedness may arise (or indeed both feelings may occur at the same time), creating a complex relationship. Similarly, gratitude can be a dispositional trait in the response to gifts, just as much as indebtedness can. We are interested in examining both dispositions and their relationship with the emergence of well-being in the context of interpersonal relationships. The measure that has been used the most to quantify the experience of indebtedness is the Revised Indebtedness Scale (IS-R, Elster et al., 2005; Van Gelder et al., 2007). This scale, based on the original Greenberg scale (1980), was designed to measure the tendency of individuals to respond to benefits received with feelings of indebtedness (feeling obligated to pay others). This measure showed a good internal consistency, and in addition, it correlated negatively with the gratitude trait.

This work aims to continue advancing with the research into the interrelationships between gratitude, debt and psychological well-being, it is of utmost importance that a measure with adequate psychometric properties be included. On the other hand, to our knowledge, there are no previous studies that have used the indebtedness trait measure with the Spanish population and with the Spanish language, so one of our first objectives will be to analyze the psychometric properties of the Indebtedness Scale (IS-R, Elster et al., 2005) in the Spanish population. The second objective will be to analyze the relationships between gratitude, debt and psychological well-being, hypothesizing that the indebtedness disposition will act as a moderator of the relationship between interpersonal gratitude and psychological well-being.

## Method

### Participants

Two independent samples were selected. A convenience sampling method was used for both cases. Participants collaborated voluntarily and received no compensation.

Fist, a sample of 229 Spanish undergraduate psychology students (162 women, 70.74%, and 67 men, 29.26%) with an age range between 18 and 45 years, M= 21.77, SD= 4.11 participated in the exploratory factor analysis, the criterion and mediational model testing. Secondly, a sample of 200 Spanish undergraduate psychology students participated in the confirmatory factor analysis (145 women, 72.5%, and 55 men, 27.5%). They depicted an age range between 18 and 45 years, M= 2180, SD= 4.45.

### Instruments

The Revised Indebtedness scale (IS-R) (Elster et al.,2005) originally developed by Greenberg (1980). This is a 22 items scale. For the adaptation of scale to Spanish, a back-translation process was performed, as previously recommended in the literature (Muñiz, Elosua, & Hambleton, 2012).The mean corrected item-total correlation was .485, Cronbach’s alpha was .88, and Spearman-Brown was .83. Thus, the revised indebtedness scale (IS-R) had good internal consistency, and might be a useful measure of this construct. Originally, the scale ranged from −3 (strongly disagree) to +3 (strongly agree), but in the current research the participants rate the items from 1 (strongly disagree) to 6 (strongly agree). This format is more familiar for participants.

Interpersonal Gratitude: Subscale of the G-20 Gratitude Questionnaire (Bernabé et al., 2014). This subscale measures gratitude for interpersonal situations. Together with the rest of the scales (gratitude to the suffering, recognition of the gifts and expression of gratitude) it seeks to measure gratitude as a construct understood in terms of existential attitude (also considering the distinction between interpersonal and transcendental gratitude). The questionnaire employs a 7-Likert scale that indicated the degree of agreement/disagreement. The scale has showed a good internal consistency, α =. 83.

*Gratitude Questionnaire-six items form* (GQ-6) (McCullough et al., 2002). In this study the spanish version used (Bernabé-Valero, García-Alandete & Gallego-Pérez, 2013) is a scale that was adapted from the GQ-6, where the item 6 was removed because of empirical and theoretical reasons. Is a 5-item self-report scale that assesses individual differences in the tendency to experience gratitude in daily life. Responses range from 1 to 7 on a 7-point Likert-type scale (1 = strongly disagree and 7 = strongly agree). Possible scores ranged from 5 to 35, with higher scores indicating a higher level of gratitude. The GQ-5 internal consistency ranged α = .77.

A number of 3 items were chosen in order to perform both convergent and divergent validity analyses. The items were the following: Q1: Relationship of Exchange: Normally, nobody gives anything without expecting something in return from you; Q2: Indebtedness and uneasiness when receiving favors; I do not like having favors done for me, because that makes me feel indebted; Q3: Happiness: Looking at your life in general, do you consider yourself to be a happy person? The items were answered using a Likert-type scale. (strongly disagree) to 7 (strongly agree).

*Ryff Scale of Psychological Well-Being* (SPWB; Ryff, 1989). A Spanish 39-item Likert scale (1 = strongly agree and 6 = strongly disagree) was used (Díaz et al., 2006). This is a self-reported scale, which assess psychological well-being from an eudaimonic conception. Individual’s well-being at a particular moment in time within each of 6 dimensions: autonomy, environmental mastery, personal growth, positive relations with others, purpose in life, self-acceptance (Ryff & Singer, 2008). For each category, a high score indicates that the respondent has a mastery of that area in his or her life. Conversely, a low score shows that the respondent struggles to feel comfortable with that particular concept. The whole SPWB showed high internal consistency α = .89, and specifically for the subscales developed in the present study, the internal consistency was: for positive relations α = .81; purpose in life α = .83 and personal growth α =. 68 (Díaz et al., 2006).

### Analysis

The analyses were developing through SPSS 22 and Amos 18.0 module. In order to examine the adequacy of indebtedness for the Spanish population in terms of psychometric properties, an exploratory factor analysis (EFA) was conducted. Assumptions were checked to ensure the application of factor analysis, such as high sample size, multivariate normality, linearity and correlation between variables (Comrey, 1973; Tabachnick & Fidell, 1989). Moreover, in order to find the suitable number of factors, Cattell’s scree-sediment graph (Cattell, 1966) as well as the eigenvalue (Kaiser, 1960). In this way, the internal consistency of the scale was evaluated through Cronbach Alpha; items of homogeneity; KMO index and the Bartlett test of Sphericity (Kaiser, 1974). After removing the factorial solution proceeded to the completion of confirmatory factor analysis (CFA) though an independent sample, accompanied by the goodness of fit indices. No rotation of the data was employed. Confirmation of the adequacy of the model have been used within the absolute fit indices; the chi-square statistic X^2^ (Jöreskog & Sörbom, 1979; Saris & Stronkhorst, 1984); and its ratio among degrees of fredoom where values under 2 are recommendable. In terms of incremental fit indices, the comparative fit index (CFI), was selected. This follows a range of values between 0 and 1 and the reference value is .90 (Bentler, 1990; Bollen, 1989; Bentler & Hu, 1998). Finally, within parsimony adjustment indices, the error of the root mean square approximation (RMSEA) of the RMSR Similarly, the more smaller its value, the better the fit, the reference value being .05 (Steiger & Lind, 1980).

Finally a mediation analysis was carried out. To conduct this analysis, the bootstrapping method of testing mediation was employed under the macro method of Hayes (2015). This method allows the measuring of the indirect effect that represents the impact of the mediator variable on the stipulated relation by a method of Bootstrapping (10000) with confidence intervals (Moret-Tatay, Lami, Oliveira, & Beneyto-Arrojo, 2018). More precisely, three variables and inherent paths were described. Fist of all, here we describe the X or interpersonal gratitude variable (independent variable), The M or indebtedness variable (mediational variable), and the Y or dependent variable (Wellbeing). A regression coefficient (and associated t-test) was calculated for each paths: on the X and M variable (path a), on the M and Y variable (path b), on the X and Y variable (c’), and in the indirect effect (whole model for path c). The logic underneath is to deepen in the construct of indebtedness and its relations with gratitude and wellbeing.

## Results

### Internal consistency

Cronbach’s alpha of the scale presented optimal values being α = .919, and the percentage of total variance explained of 49.17%. Table 1 presents the descriptive analysis, homogeneity items, Cronbach’s alpha, kurtosis, skewness and exploratory factor loadings between items.

**Table 1.** Means, standard deviation, item homogeneity, kurtosis, skewness and exploratory factor loadings for the four items of the

| Item wording | Mean | SD | H<sup>2</sup> | Kurtosis | Skewness | Cronbach's $\alpha$ |
| --- | --- | --- | --- | --- | --- | --- |
| Factor 1 | 4.08 | 1.05 | .41-.65 | -.32 | -.44 | .81 |
| Factor 2 | 2.76 | 1.05 | .44-.64 | -.47 | .33 | .82 |
| Factor 3 | 3.51 | 1.08 | .54-.80 | -.50 | .06 | .83 |
| Factor 4 | 2.93 | 1.15 | .42-.64 | -.53 | .36 | .75 |

SD= standard deviation; H<sup>2</sup>= Homogeneity range

### Exploratory and Confirmatory Factor Analysis

In relation to the validity of Exploratory Factor Analysis (EFA), the Bartlett’s test of sphericity was p <.001 with a value of chi-square 2254.78 (df = 231) and the sample index value of Kaiser-Meyer-Olkin (KMO) was 0.93. The scree-test (Cattel, 1966) recommended four factor solution. The AFC has confirmed through and independent sample (n=200) that the existence of four factors. The model presented an optimal fit. The goodness of fit indices global scale was: X^2^ = 363.21 p <.001 (df = 203), X^2^ /df=1.79, CFI = .914, IFI = .915, and RMSEA = .06.

### Validity

In terms of qualitative analysis of the factors from the items, it has been pointed out that each factor shows a high coherence in terms of content. Moreover, with regards to the measurement of different facets of the debt appearing on the scale. In order to summarize the information of each factor, factors were labeled as follows:

Factor 1: Debt for material aspects (Items 3,5,6,7,8,9 and 12): in this factor the information related to the discomfort or aversion to contract material debts is described.
Factor 2: Self-sufficiency and discomfort in receiving help (Items 2, 4, 11, 13, 14 and 15): refers to the discomfort of receiving help and favors from others with an attitude of self-sufficiency
Factor 3: Moral self-demand in the reception of help (Items 1, 10, 18, 19 and 21): a high score in these items indicates that the participants go beyond the rules of exchange; they are self-demanding to be quick in returning the favor. Moreover, the feeling of obligation to return more than the favor received is described here, as well as the return of all favors as indicative of being “good friend”.
Factor 4: Debt in the receipt of gifts (Items 16, 17, 20 and 22) describes various emotional responses of discomfort, concern and not enjoyment when receiving gifts and relief if they are not received.

To test the criterion validity of the scale, this was correlated between the gratitude and happiness construct and other theoretical constructs associated with debt. More precisely, with items on the Exchange Relationships (Q1), the discomfort with the reception of favors (Q2) and Happiness. Table 2 depicts the positive correlation of debt with Q1 and Q2. As expected, Debt did not correlated with Happiness (except for the Factor 4) or Gratitude (except for Factors 2 and 4). Finally, figure 1 shows the final factor structure, in terms of factor loading.

**Table 2.** Pearson coefficients among Factors R-Indebtedness, Q1, Q2 and Happiness for construct validity

|  | Q1 (Debt) | Q2 (Debt) | Happiness | Gratitude | Factor 1 | Factor 2 | Factor 3 | Factor 4 |
| --- | --- | --- | --- | --- | --- | --- | --- | --- |
| Q1 (Debt) | 1 |  |  |  |  |  |  |  |
| Q2 (Debt) | **.439\*\*** | 1 |  |  |  |  |  |  |
| Happiness | -.122 | **-.136\*** | 1 |  |  |  |  |  |
| Gratitude | **-.212\*\*** | **-.233\*\*** | **.489\*\*** | 1 |  |  |  |  |
| Factor 1 | **.207\*\*** | **.349\*\*** | -.032 | -.112 | 1 |  |  |  |
| Factor 2 | **.340\*\*** | **.470\*\*** | -.113 | **-.200\*\*** | **.636\*\*** | 1 |  |  |
| Factor 3 | **.260\*\*** | **.243\*\*** | -.050 | -.100 | **.626\*\*** | **.652\*\*** | 1 |  |
| Factor 4 | **.264\*\*** | **.289\*\*** | **-.155\*** | **-.172\*\*** | **.445\*\*** | **.572\*\*** | **.594\*\*** | 1 |

\* $p < .05$ ; \*\* $p < .01$

**Figure 1.**
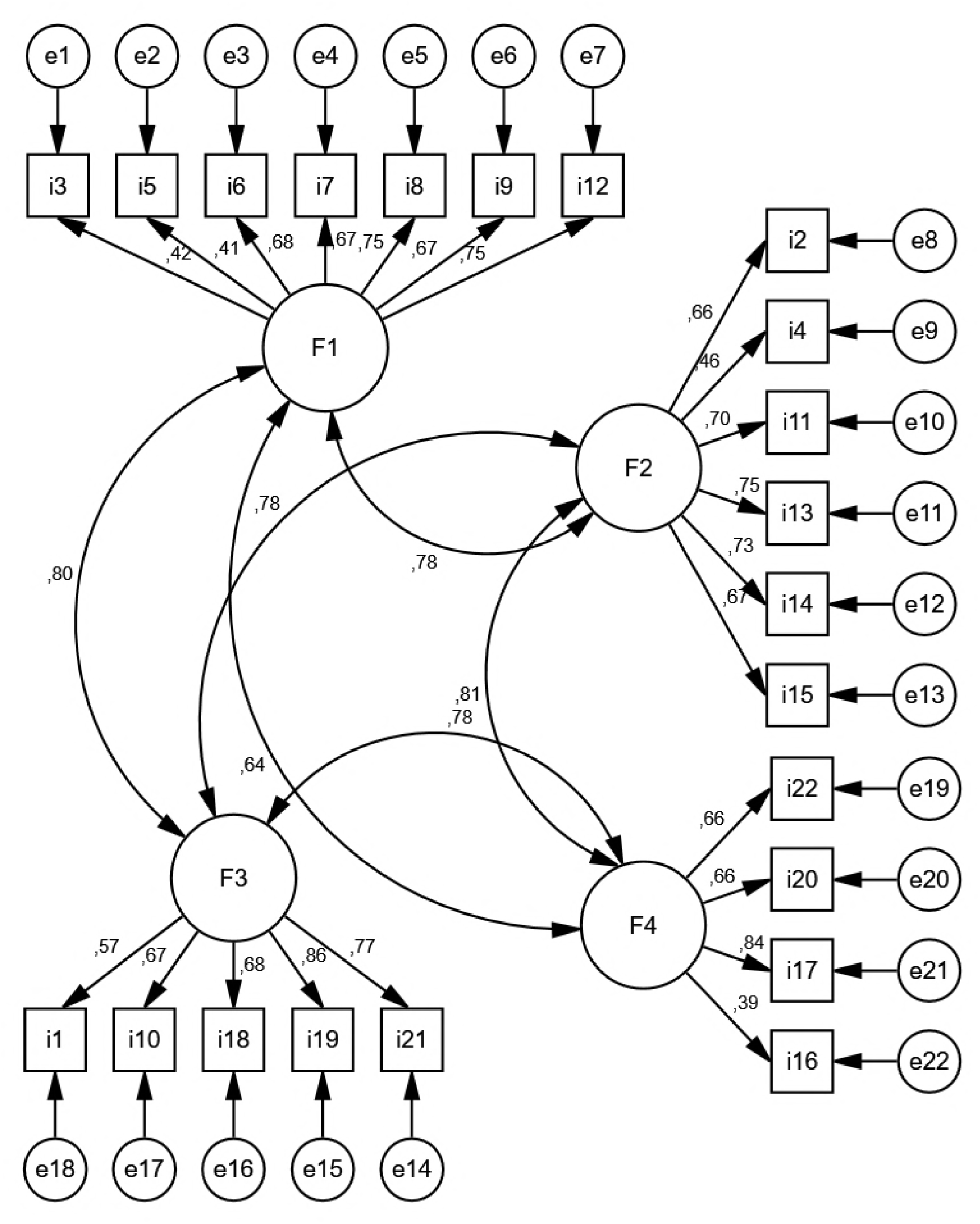
Factor loadings and structure for the debt scale

A mediational model to test indirect effects (see figure 2). This analysis also allows us to determine whether indebtedness mediated the relation among interpersonal gratitude and Wellbeing. Thus, this was carried out through interpersonal gratitude, the different subfactors of indebedtness and wellbeing. Only subfactors for Self-sufficiency and discomfort in receiving help and positive relation condition fulfilled all assumptions for this analysis (see figure 2). The overall model was statistically significant: F_(1,227)_=4.98, p<.05, R^2^=.15. In terms of path, first, the path of X and M (path a) was tested, which resulted in a positive significant effect (β = -.15, t=2.23, SE=.07, p < .05). The second path b, among M and Y, was tested, reaching the statistical level: (β = -.37, t=6.97, SE=.05, p <.0001). The c’ also reach the statistical significance: (β = .17, t=2.98, SE=.05, p <.05). The total effect reached the statistical significance: (β = .22, t=3.67, SE=.05, p <.0001). Following Kenny and Baron (1986), the causal method steps are satisfied as path b> c’ and c’<c.

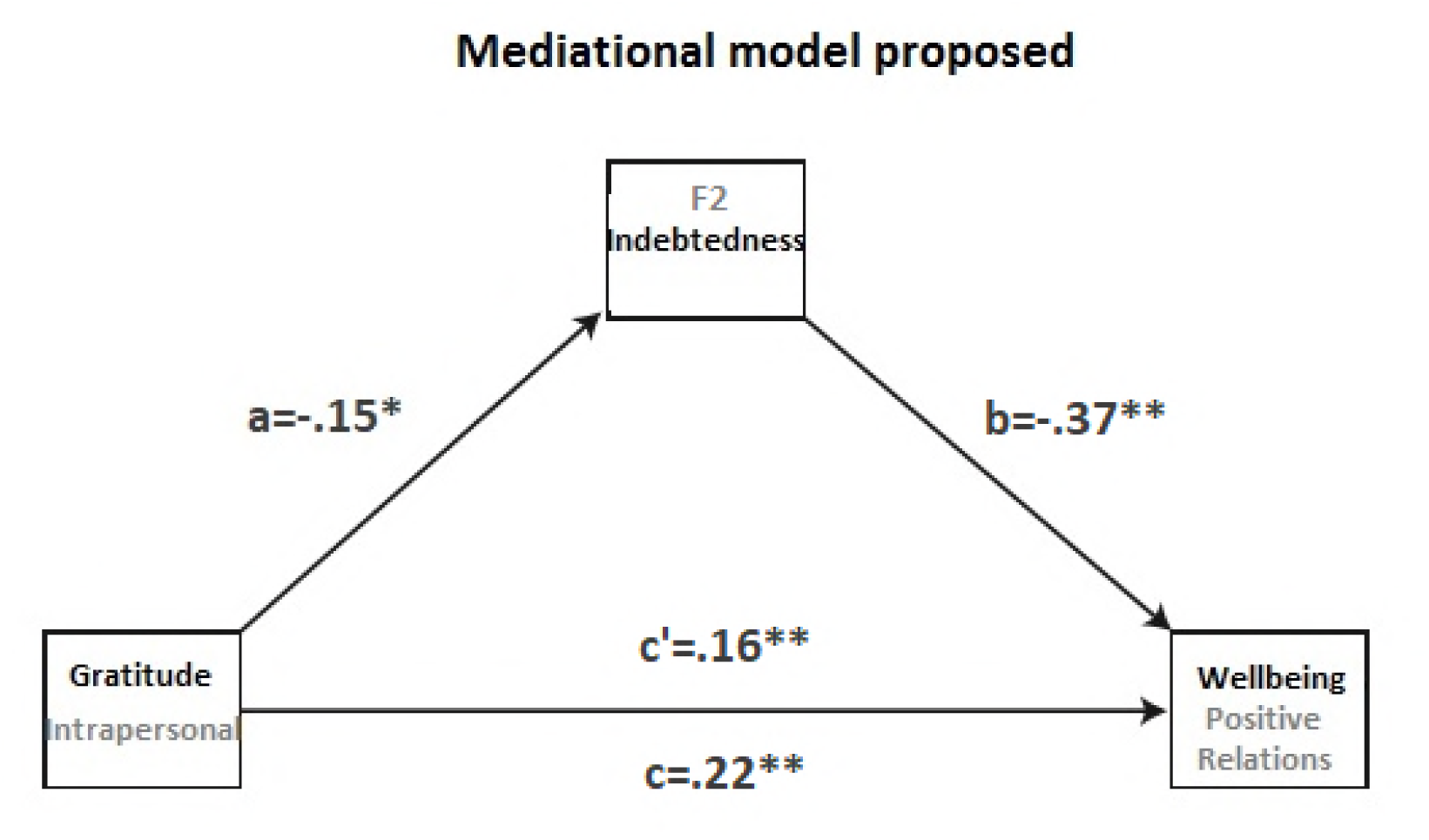
Figura 2. Mediational model proposed across indebtedness, interpersonal gratitude and Wellbeing.

## Conclusions and discussion

The aim of this work was to examine the role of indebtedness in well-being. For this reason, the R-Indebtedness Scale was adapted to the Spanish population and explored in terms of psychometric properties. A 4-factor solution structure was found with interrelated factors. The factors have a high coherence in their content, so it has been possible to label and describe them.

With regards to the convergent validity, we can affirm that it is high since there are significant, positive correlations between the 4 factors of the R-Indebtedness scale and the Q1 of Exchange Relationships and the Q2 of Discomfort upon the receipt of favors. With regards to discriminant validity, we observed that there is no correlation between Q3 (Happiness) and F1, F2 and F3, supporting the idea of good discriminant validity. It could be the case that F4 (Debt in the receipt of gifts) involved starting from an underlying mindset (dislike for receiving gifts and relief if they are not received), which was inversely related to happiness, although this would not be the same construct.

On the contrary, the relationships between gratitude and indebtedness are more complex: on the one hand, all the indebtedness factors are inversely correlated with gratitude as the GQ5, although only F2 (Self-sufficiency and discomfort in receiving help) and F4 (Debt in the receipt of gifts) are so to a significant extent.

We hypothesize that F1 (Debt for material aspects) and F3 (Moral self-demand in the reception of help) imply less aversion, and therefore, aspects of indebtedness compatible with gratitude are conceivable and they may not necessarily have an inverse relationship. In other words, people who scored high in these two aspects of indebtedness may feel the importance of returning the favor and not wanting to owe material aspects, but this would not always be related to the fact of being able to thank the positive aspects of the reception of the gift. These dispositional aspects of indebtedness would not produce a decrease in the disposition to gratitude, and may be at the expense of other situational factors such as the emergence of the gratitude emotion. Therefore, we propose that there are various degrees of negative emotionality surrounding indebtedness, and that factors 2 and 4 may have a greater degree of discomfort or negative emotionality. More research would be required in this area.

On the other hand, these results are consistent with the mediating model we have tested, in which it is indicated that only the model that included the F2 could have a good fit. With regards to the mediational model, different models were tested with the different factors of the indebtedness scale (IS-R, Elster et al., 2005), as well as including the GQ-5 (McCullought et al., 2002) and the Interpersonal Gratitude (G-20, Bernabé et al., 2014) measurements. In turn, models were tested which included the different dimensions of Psychological Well-Being (SPWB; Ryff, 1989), self-acceptance, positive relations with others, autonomy, environmental mastery, purpose in life and personal growth. Only the model that included the Factor 2 of indebtedness, interpersonal gratitude and positive relationships with others (Ryff) obtained a good fit. As expected, the “Self-sufficiency and discomfort in receiving help” measured the relationship between interpersonal gratitude and positive relationships.

This result is also consistent with the proposals that indicate that self-sufficiency is negatively related to gratitude (for example, Buck, 2004). It would be interesting to research the mental representations of the others (“internal working model of others”) and its relation to the disposition to indebtedness and gratitude. For instance, Mikulincer and Shaver (2010) showed how avoidant attachment, with its deactivating strategies of the attachment system (e.g., seeking emotional distance and self-sufficiency) exacerbated the negative views of others by diverting attention from relevant information for attachment (information about positive traits, intentions and actions of others), leading to less gratitude and interfering with the prosocial response. We hypothesize that including the attachment style in the studies on the relationship between gratitude, indebtedness and psychological well-being would shed more light on this area. The results of this model continue to confirm the positive relationship that exists between gratitude and personal relationships (Algoe, 2012, Gordon, Impett, EKogan, Oveis & Keltner, 2012, Kubacka, Finkenauer, Rusbult, & Keijsers, 2011). For instance, in a recent study (Algoe & Zhaoyang, 2016) it was shown through an experiment, that enhancing gratitude in close relationships improved personal and relational well-being, in comparison with having another type of interaction that is believed to generate more relational intimacy. Therefore, from our results, we propose that it would be interesting to include in these types of studies the disposition to indebtedness (in all its different facets) to be able to define this relationship to a greater extent and to know the key elements that may hinder the development of social ties and progress in personal well-being. In addition, being aware of these elements can be an important factor in the design and handling of a therapeutic work designed to increase gratitude in people. Moroever, these results highlight the importance of having conceptually well-defined instruments with good psychometric properties, such as the IS-R scale that we have analyzed. The fact of being able to count on the 4 labeled factors allows us to carry out different analyzes with greater precision. In addition, it is a useful tool to be used in various statistical analyzes such as Mediation and SEM. In the same way, we have been able to benefit from an instrument of measurement that defines the types of gratitude (interpersonal, transcendental and faced with suffering), and with another one which defines the different facets of well-being, which has brought more conceptual coherence to the empirical analyzes.

These results are of interest not only for an applied level, it also for a theoretical one. One should bear in mind the relation between gratitude and well-being. A review of studies using positive psychology techniques (Sutipan, Intarakamhang, & Macaskill, 2017) suggested that such techniques serve as possible useful tools for the future to improve well-being, happiness, life satisfaction, and relieve depressive symptoms in the elderly. From all these studies, the relationship between gratitude and well-being appears to be proven. However, we agree with Nezlek, Newman & Thrash (2017) when they state that, although there is a significant relationship between gratitude and well-being, these relationships vary depending on how well-being is defined. In the current work, we argue that these relationships may also vary depending on how gratitude is defined. Thus, gratitude has been defined and investigated from several conceptualizations (such as emotion, affective trait, existential attitude, virtue among others.), as well as with various measuring instruments (eg Bernabé, García & Gallego, 2014; McCullough et al., 2002; Watkins, et al., 2003; Wood, Maltby, Stewart, & Joseph, 2008). All the definitions seem to agree on the fact that one of the privileged areas in which this construct manifests is in interpersonal relationships. It is in this context that people can offer gifts and help and therefore, they can feel and express gratitude. The explanatory model on which we are based (Bernabé, 2014) considers that gratitude could be defined as a predisposition of the person to recognize, value and respond to the positive aspects of personal existence, experienced as gifts received. Moreover, it makes a distinction between transcendental gratitude (towards God or existence) and interpersonal gratitude, depending on the agent towards whom gratitude is felt. It also differentiates types of gratitude according to the object, proposing, in addition, gratitude related to suffering. These distinctions between the types of gratitude are especially important in order to deepen the relationship between gratitude and well-being. We argue that interpersonal gratitude is more directly related to the promotion of social ties and prosociality, since it is what emerges in the context of the exchange of gifts or favors, and is what enables the strengthening of personal relationships. At the same time, transcendental gratitude might further promote personal (eudaimonic) wellbeing through the assessment of the positive aspects of personal experience. However, we understand that both would be related since, the increase and maintenance of the social bonds created by gratitude, would be a powerful mechanism that would explain the relationship between gratitude, well-being and health (McCullough, Kimeldorf, & Cohen, 2008). In fact, one of the subscales of a very important instrument of psychological well-being, the Ryff Scale of Psychological Well-Being (SPWB; Ryff, 1989), refers to positive relationships. This instrument that comprises six distinct dimensions including self-acceptance, positive relations with others, autonomy, environmental mastery, purpose in life and personal growth. These dimensions together can contribute to the appraisal of positive functioning and well-being. Within this relationship between gratitude and well-being, those aspects that may interfere in the response of gratitude when receiving a benefit provided by other people seem of interest (considering it essential to know those variables that can play an inhibitory role in the emergence of gratitude). In the previous literature, we have also found the following variables inversely related to gratitude; cynicism, materialism, envy, narcissism and indebtedness. (Solom et al., 2017). Some of them might have a closer relationship with the emergence of interpersonal gratitude and others with transcendental gratitude. For example, regarding materialism and envy, we find different cross-sectional studies that affirm that these two variables are negatively associated with the gratitude trait (Kashdan & Breen, 2007; Lambert, Fincham, Stillman & Dean, 2009; McCullough et al., 2002; Polak & McCullough, 2006). The results suggest that materialism implies that people wish to obtain more and more possessions and, therefore, do not value the things they already have in their lives. Consequently, this attitude does not necessarily relate directly to the emergence of gratitude in the context of an interpersonal relationship. In the same way, envy could arise from an ascending social comparison in which people may consider that others have more or better positive aspects in their existence.

